# Event-driven acquisition for content-enriched microscopy

**DOI:** 10.1101/2021.10.04.463102

**Authors:** Dora Mahecic, Willi L. Stepp, Chen Zhang, Juliette Griffié, Martin Weigert, Suliana Manley

## Abstract

A common goal of fluorescence microscopy is to collect data on specific biological events. Yet, the event-specific content that can be collected from a sample is limited, especially for rare or stochastic processes. This is due in part to photobleaching and phototoxicity, which constrain imaging speed and duration. We developed an event-driven acquisition (EDA) framework, in which neural network-based recognition of specific biological events triggers real-time control in an instant structured illumination microscope (iSIM). Our setup adapts acquisitions on-the-fly by switching between a slow imaging rate while detecting the onset of events, and a fast imaging rate during their progression. Thus, we capture mitochondrial and bacterial divisions at imaging rates that match their dynamic timescales, while extending overall imaging durations. Because EDA allows the microscope to respond specifically to complex biological events, it acquires data enriched in relevant content.

## Introduction

Live-cell fluorescence microscopy is an indispensable tool for studying the dynamics of biological systems. Yet, fluorescence microscopy also comes with limitations: as fluorophores go through excitation and emission cycles, they photobleach and produce reactive oxygen species, which interact with cellular components, causing phototoxicity. Thus, photobleaching and phototoxicity restrict the photon budget. Practically, this manifests as a trade-off between spatial resolution, temporal resolution, measurement duration, and signal-to-noise ratio (SNR), such that improving one parameter comes at a cost to the others^1,2^. For example, higher imaging rates degrade photon budget and sample health more rapidly, reducing measurement duration and SNR. Super-resolution microscopies resolve features below the diffraction limit, but also require higher light doses.

Methodological developments have addressed the limited photon budget with several complementary approaches. Brighter and more photostable fluorescent probes increase the photon budget^3,4^, while computational denoising or reconstruction^5–8^ restores low SNR or sparse images to use photons more efficiently, and adaptive microscopies improve how measurements allocate photons. These approaches may be combined synergistically, to permit access to longer, gentler measurements.

Adaptive microscopies implement feedback control systems that spatially modulate the excitation and reduce the overall light dose. Modulated scanning can adjust laser dwell times or intensities to amplify pixels with higher fluorescence signals, to enable faster and gentler confocal^9,10^ and super-resolution imaging^11,12^. Alternatively, modulation can maintain pixel-wise fluorescence intensities, and prioritize increased dynamic range while maintaining light exposure^13^. Additional inputs, such as prior knowledge of sample organization or shape have been used to optimize local excitation intensity in multi-photon measurements^14,15^. In widefield, digital micro-mirror devices can illuminate selectively where fluorescent structures reside, and reduce the overall light dose^16^. These adaptive approaches generally take into account the spatial heterogeneity of biological samples, and use their fluorescence signal to decide where, and how much to illuminate.

Biological processes, however, also occur over a broad range of temporal scales and with complex dynamic signatures. For example, within eukaryotic cells, mitochondrial fissions are sporadic events which typically only occur once every few minutes in an entire mammalian cell, and persist less than a minute from start to finish^17^. The constriction rate accelerates towards full scission – the final stages of which contain the richest information about membrane deformation and its mechanisms. By contrast, the bacterial cell cycle is deterministic, but also intermittent; cells elongate and partially constrict, before popping apart into two separate daughter cells during a small fraction of the cell cycle duration (0.1-0.0001%)^18,19^. As a result, such dynamic biological events are challenging to capture, especially under their native conditions without induction. Measurements taken at a fixed frame rate would risk missing events of interest, since a high frame rate limits measurement duration while a slow frame rate limits event-specific content. Thus, the optimal frame rate, set by the biological dynamics, changes during an acquisition.

Open-source control software enables users to define and execute their own automation settings, adapted to their biological samples and for their own instruments (MicroManager Intelligent Acquisitions plugin^20, 21,22^). However, biological samples often undergo subtle changes in morphology and protein assembly patterns that herald the onset of events of interest. Under these conditions, adaptive control based on simple readouts like intensity^23^ or cell shape^22^ would lack the required sensitivity and specificity to detect them. The complexity of this challenge has prevented event-driven control from achieving its full potential to enable biological discovery through extension beyond spatial adaptation to include temporal adaptation, sculpting the measurement to enrich for events of interest.

We introduce an event-driven acquisition (EDA) framework, which responds to the distinct spatial signatures of biological events to adapt the microscope acquisition – as demonstrated here, by modulating the frame rate. Image analysis powered by deep learning is capable of recognizing surprisingly subtle signatures of events of interest within the sample. We harness a computer vision approach by training a neural network to detect precursors to events of interest and tailor the acquisition on-the-fly. To aid implementation of EDA in different microscope setups, we provide a plugin that integrates the neural network analysis and actuation into Micro-Manager. As proof of principle, we integrate EDA into a large field-of-view instant structured illumination microscope (iSIM)^24,25^ and apply it to capture super-resolved time-lapse movies of mitochondrial and bacterial divisions. In response to neural network outputs, EDA prioritizes the imaging speed or measurement duration. Such acquisitions spend the sample’s photon budget more efficiently compared with a fixed frame rate. Controlled by EDA, the iSIM collects data enriched in event-specific content and captures more completely the dynamic progression from constriction through scission.

## Results

### EDA concept and implementation

#### Light Exposure Imaging Speed

The EDA framework consists of a feedback loop between the live image stream and microscope controls, whose implementation requires: 1) sensing via images of the sample collected by the microscope, 2) computation to analyze images and detect events of interest and 3) actuation to adapt the measurement parameters (Fig. 1a). In our implementation, the open software MicroManager^20^ captures images from the microscope. As new frames are received, a neural network trained on labeled bio-image data to recognize specific events of interests performs the computation step. For each image, the network generates a spatial relative probability map, or ‘event score’, for those events of interest. The event score acts as a decision parameter for the control loop, and actuates changes in the measurement. We offer this framework as a Micro-Manager plugin – capable of multithreading to parallelize image analysis – that allows EDA-enabled experiments on a wide variety of microscopes (Supplementary Note 1). We demonstrate its utility here for measurements of mitochondrial and bacterial division events taken with our custom-built iSIM (Methods).

**Fig. 1:**
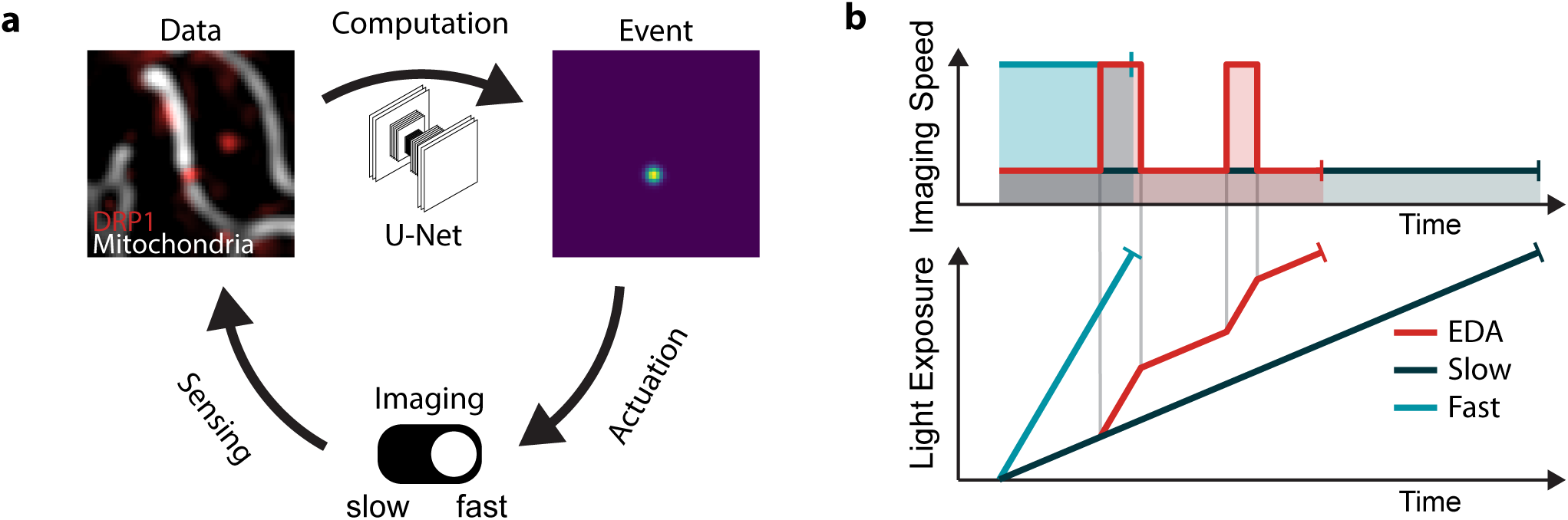
Event driven acquisition (EDA) concept. **a**, The feedback control loop for EDA is composed of three main parts: 1) sensing by image capture to gather data, 2) computation by a neural network to detect events of interest and generate a heat map of event score, and 3) adaptation of the acquisition parameters in response to events in the sample. **b**, Schema of experiment duration and cumulative light exposure for different fixed imaging speeds or with EDA to switch between the two. The total excitation light dose (shaded areas in upper panel, final light exposure value in lower panel) is constant for all experimental configurations to represent the fixed photon budget.

Since our events of interest are intermittent, we use the event score to actuate a toggle between slow and fast imaging speeds as the measurement proceeds (top, Fig. 1b). We rely on the global maximum event score found in the image, reflecting the likelihood of event-relevant content. When it exceeds a user-defined value, the microscope begins to acquire data at a high frame rate. Once in this fast mode, the microscope switches back to the slow mode only when the event score diminishes below a second, slightly lower, threshold. This hysteresis in the triggering mechanism makes the actuation less susceptible to noise from the readout and ensures that events are followed for their full duration, even if the event score peaks early during the event. Given a limited photon budget and fixed frame rate, the measurement will last longest for the slow imaging speed and shortest for the fast imaging speed. The EDA measurement has an intermediate duration, but spends more of its photon budget during events of interest, when the sample will accumulate more exposure to light (bottom, Fig. 1b).

### Detecting division using a neural network

Division underlies mitochondrial proliferation and maintenance^26^, similar to its role in free-living single-celled organisms. The structural and dynamic precursors to mitochondrial division are challenging to detect. They can occur infrequently, and at any location within the mitochondrial network, at any time. Mitochondria are also constantly in motion, and undergo shape changes unrelated to division. Once initiated, constrictions typically proceed to division within less than a minute^17^. Finally, mitochondrial width lies near the diffraction limit; thus, changes in their shape preceding fission are difficult to measure. Although numerous software packages propose automated segmentation^27,28^ and tracking of mitochondria as well as network morphology classification^29–32^, they rarely attempt to detect divisions^33^ or active constrictions preceding them. For these reasons, microscopists typically rely on manual analysis to identify mitochondrial divisions in their data, playing videos forward and backward and visually inspecting local changes in the network. Overall, detecting division events in real-time during imaging presents a major bottleneck.

The dynamin-related protein 1 (DRP1) drives mitochondrial membrane constriction and scission, and is required for spontaneous divisions^34–36^. Thus, its enrichment at division sites can help to identify them. But DRP1 also accumulates at nondividing sites, since it can bind and oligomerize on both curved and flat membranes^37^. So, its presence alone cannot reliably predict future divisions, precluding a simple intensity threshold-based event detection.

To address these challenges and detect mitochondrial division sites, we rely on the combined information from mitochondrial shape and DRP1 enrichment. We trained a neural network with a U-net architecture^38^ on input images of mitochondria and DRP1. The shape of a constriction site is a saddle point, which is characterized by two principal curvatures of opposite signs.

Those curvatures are represented by the eigenvalues of the Hessian matrix at that point in the image. Thus, as a putative ground truth, we used the intensity in the DRP1 channel, multiplied by the negative determinant of the Hessian matrix representing the Gaussian curvature of the mitochondrial outline (Supplementary Note 2, Methods). Importantly, to generate the final ground truth data, we manually curated the training set to increase the accuracy for real constriction events by removing false positives.

Our network, trained to detect mitochondrial constrictions in the presence of DRP1 (Fig. 2a), produces a heat map of event scores: higher values mark locations within the image where division is more likely to occur ((Fig. 2b), Supplementary Figure 4). The maximum event score represents the most pronounced constriction site highly enriched in DRP1, and hence the most relevant content in a given frame. This network output acts as a decision-making parameter for the actuation step in the EDA feedback control loop (Fig. 2c). Once the maximum event score exceeds a threshold value, the imaging speed increases after the subsequent frame, and remains high until the event score dips below a second value. In the example shown, the neural network detects two events of interest, which trigger fast imaging in the iSIM; the first constriction event relaxes without dividing, while the second one ends in division.

**Fig. 2:**
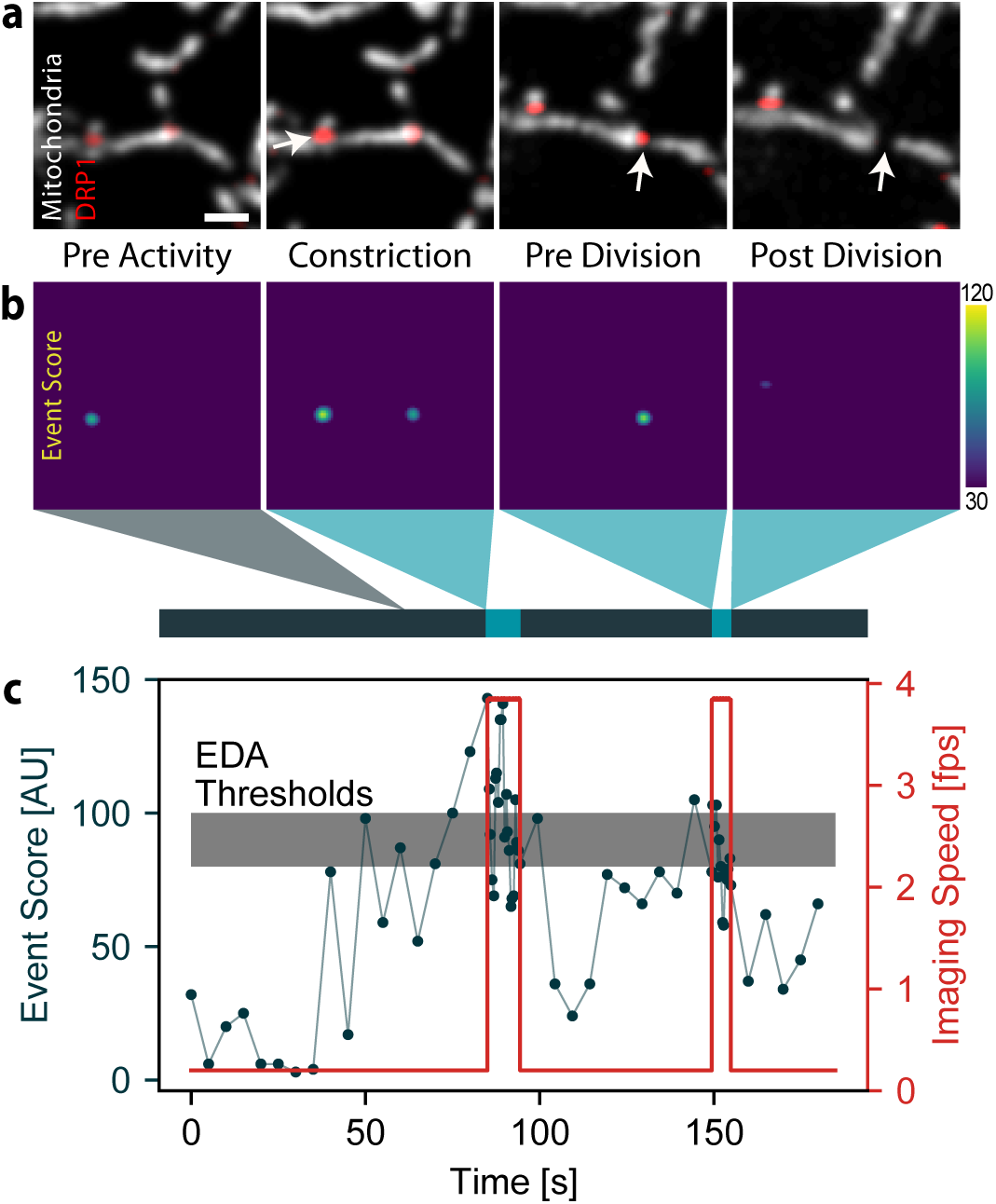
Event recognition of mitochondrial divisions during an iSIM acquisition. **a**, Images of a COS-7 cell expressing mitochondrion-targeted Mito-TagRFP and Emerald-DRP1, including those containing events of interest (white arrow) that triggered a change in the imaging speed. (Scale bar: 1 µm; image intensities were corrected for photobleaching) **b**, Corresponding event score maps and measurement timeline (grey, slow speed; blue, fast speed). **c**, Time course of the maximum event score computed for each frame of the acquisition (black), and the adaptive imaging speed actuated by the event score (red). The upper boundary of the grey band denotes the maximum threshold above which the iSIM switches to fast imaging, and the lower boundary the minumum threshold below which it returns to slow imaging.

### Temporally adaptive imaging of mitochondrial division

We performed EDA with our iSIM on COS-7 cells expressing mitochondrion-targeted Mito-TagRFP (Cox-8 presequence) and Emerald-DRP1. Typically, our FOV included one entire cell, and we focused in a plane near the coverslip, where many mitochondria were visible. During each measurement, the network recognized events of interest which triggered a switch from slow to fast imaging, followed by a return to slow imaging. Many sites that accumulated DRP1 did not lead to division, and also failed to trigger the network (Fig. 3a), demonstrating the network’s ability to discriminate events of interest.

**Fig. 3:**
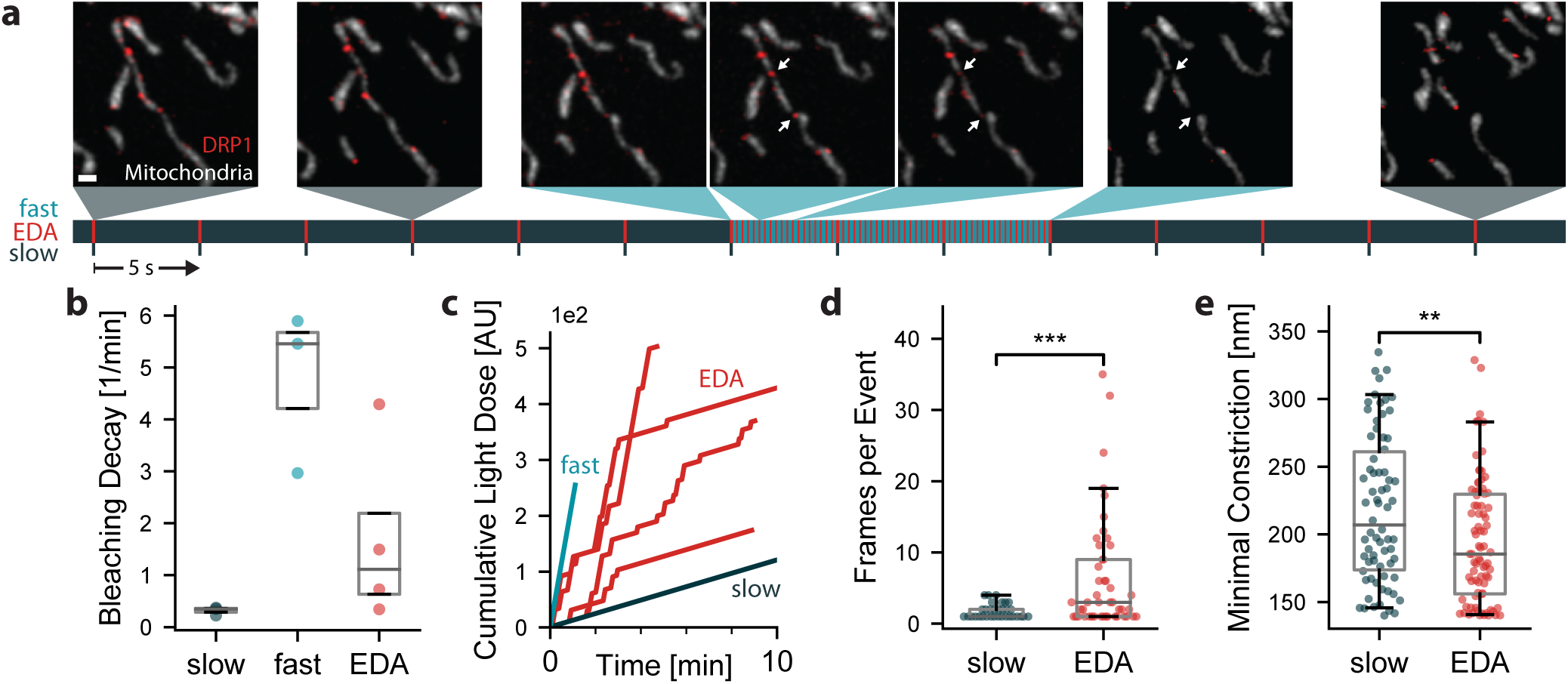
EDA versus fixed-rate imaging of mitochondrial divisions. **a**, Mitochondrial dynamics (TagRFP-Mito, grey; Emerald-DRP1, red) captured by EDA (Scale bar: 1 µm, time first frame to last frame: 114 s). Below, measurement timeline indicating the approximate capture time of each image (grey, slow speed; blue, fast speed; red, frame capture). **b**, Decay constants obtained from exponential fits to the time-dependent mean intensity in the mitochondria (Supplementary Figure 1). Each point corresponds to a separate FOV. **c**, Cumulative light dose, defined as the laser power multiplied by the exposure time summed over all previous frames. EDA sometimes achieves a higher total light dose than fast imaging alone, due to recovery during slow imaging (Supplementary Figure 1). **d**, Number of frames with a constriction diameter below 200 nm during the 20 seconds after the event score exceeded the threshold value. **e**, Minimal constriction width measured during the frames in d. (slow: N = 360 frames in n = 3 independent experiments; fast: N = 765, n = 2; EDA: N = 1516, n = 4; Box plots mark the first quartile, median and third quartile with the whiskers spanning the 95 % confidence interval. Significance tests were performed using the independent two-sample t-test.

To compare the duration of EDA to traditional imaging acquisitions using fixed parameters, we also collected data at a constant imaging speed of either 0.2 frames/sec (slow) or 3.8 frames/sec (fast). EDA showed less photobleaching decay relative to fast imaging, with an average 3-fold decrease in the exponential bleaching decay constant (Fig. 3b). While the bleaching decay is still on average 5 times higher relative to slow imaging alone, it was possible to reach 10-minute imaging durations for many EDA experiments without excessive photobleaching or sample health degradation. EDA measurements sometimes showed photobleaching recovery while imaging at a slow speed (Supplementary Figure 1), enabling a higher cumulative light dose compared with fixed-rate fast imaging (Fig. 3c). During EDA measurements, events of interest toggled the actuation from slow to fast on average 9 times for an average of 10 seconds resulting in EDA imaging fast for 18 % of the frames.

The potential of intelligent microscopy includes measuring what standard acquisitions would miss – beyond measuring for longer overall durations. Ideally, EDA should collect constriction states leading up to division that would be inaccessible to slow imaging. We therefore use constriction diameters to quantify fission intermediates as a measure of event-relevant content, independent of the event score. During division events, (Fig. 2c), we asked how many frames EDA collected with highly constricted diameters (below 200 nm) compared to slow fixed-rate imaging (Fig. 3d). The resolution of raw iSIM images is 210 nm prior to deconvolution. Therefore, information about constriction diameters below 200 nm is unavailable during on-the-fly processing, and reflects the ability of EDA to collect event-relevant content distinct from the ground truth for the neural network. We found the average number of such frames (constriction < 200 nm) during the first 20 seconds of an event was 5.4-fold higher for EDA compared with fixed slow imaging (7.5 versus 1.4 frames). This content enrichment is also reflected in the distribution of minimal constriction diameters (Fig. 3e), both in terms of its lower mean value and its skew. For fixed fast imaging, a similar amount of data only showed three constriction events (and thus is not included in the figure panel), further highlighting the limitations of fixed-rate imaging compared with EDA. Thus, EDA better resolves the constricted states preceding division, as well as the progression of molecular and membrane states leading to fission, captured by each burst of fast images (Supplementary Figure 2).

### Extending EDA to bacterial cell division

An endosymbiotic alphaproteobacterium is believed to be the ancestor to mitochondria^39^. Over time, the proteins required for division were replaced by homologues encoded in the host genome, which assemble on the mitochondrial exterior as opposed to the bacterial interior. A superficial resemblance remains between the sub-micrometric diameters of mitochondria and many bacteria, and the shape transitions they both undergo en route to division. But, the bacterial cell cycle occurs on the timescale of tens of minutes – over which the cells elongate and partially constrict, before popping apart into two separate daughter cells during a small fraction of the cell cycle duration (0.1-0.0001%)^18,19^ – in contrast with the tens of seconds for mitochondrial division. Thus, it poses a distinct set of challenges to live-cell microscopy.

We extended EDA to study the final stages of division of the alphaproteobacterium *Caulobacter crescentus*. We imaged cells expressing the cytoplasmic fluorescent protein mScarlet-I, and the bacterial divisome protein FtsZ-sfGFP (Fig. 4a, Supplementary Note 3) with the iSIM. In newborn *C. crescentus*, FtsZ forms a punctum associated with one cell pole. As cells elongate, FtsZ depolymerizes and appears diffuse, before assembling into a narrow band of filaments that coordinates constriction at midcell. The band shrinks as the cell constricts, and finishes as a punctum at the division site. We found that our event detection network developed for mitochondrial constrictions could recognize the final stages of *C. crescentus* division without any additional training (Supplementary Note 2). This transferability is likely due to morphological similarities in constriction shape and the presence of a functionally similar molecular marker. High event scores remained specific to states close to division, preventing actuation from being triggered by the punctate polar arrangement of FtsZ at the beginning of the cell cycle (G0). *C. crescentus* cells at all stages of cell cycle were plated onto an agarose pad (Methods), with several hundred in the iSIM FOV. We collected data on cells at a fixed slow imaging speed of 6.7 frames/hour, a fast imaging speed of 20 frames/hour, or a variable speed switched by EDA. The bleaching decay rate diminished compared with the fixed fast imaging rate by a factor of 1.7 (Fig. 4b). As for mitochondria, we found that constrictions measured with EDA had significantly smaller diameters on average compared with fixed slow imaging (Fig. 4c), with the overall distribution shifted to lower values. This confirms that EDA improves access to details of bacterial cell division that are difficult to capture using a fixed imaging speed, and uses the photon budget more efficiently.

**Fig. 4:**
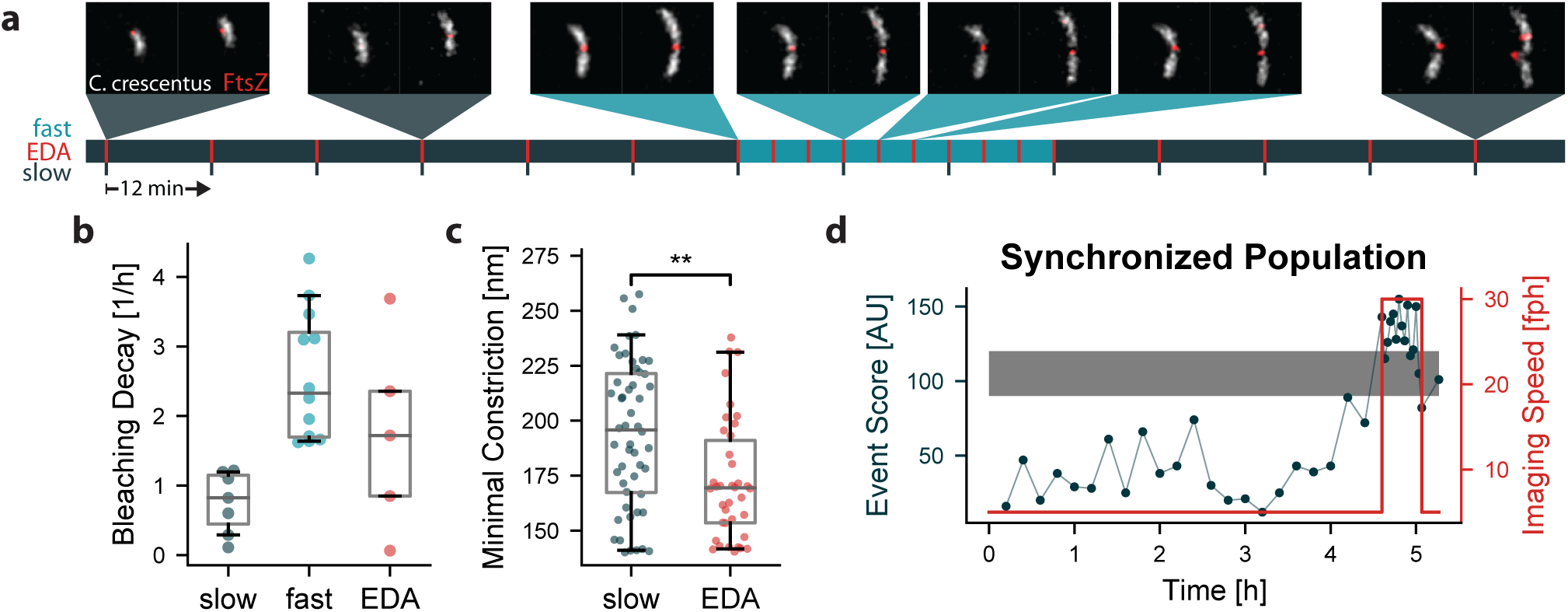
EDA versus fixed-rate imaging of bacterial divisions. **a**, Representative images of *C. crescentus* (cytosolic mScarlet-I in grey and FtsZ-sfGFP in red) division events captured at different imaging speeds using an event-driven acquisition (Scale bar: 1 µm, time first frame to last frame: 5.8 h). **b**, Decay constants obtained from exponential fitting of the mean intensity in the bacteria over time. **c**, Minimal constriction width measured when following a constriction event over time. **d** Event score and imaging speed for an EDA triggered experiment on cell-cycle synchronized bacteria. Long lag phase and many divisions at the same time imaged in high temporal resolution triggered by EDA. (slow: N = 228, n = 2; fast: N = 296, n = 3; EDA: N = 182, n = 5).

We also used EDA to image bacterial populations that were first synchronized into the beginning of the swarmer (G0) phase of cell cycle through density centrifugation (Methods). Initially, EDA maintained a slow imaging speed during the lag phase while bacteria increased in length without dividing (Fig. 4d). This allowed a low light dose, so that cells could develop without division arrest and filamentation which occurred at faster imaging speeds. After several hours, the first wave of divisions in the population occurred, and EDA triggered high-speed imaging. In this manner, EDA captured many divisions simultaneously, as a consequence of a population-wide high and stable event score. After the population fully divided, imaging went back to the slow imaging speed, preserving the photon budget for the next wave of divisions (Supplementary Figure 3). Synchrony allowed for a further boost in events of interest, with EDA sensing and responding with the suitable timing based on cell-cycle progression.

## Discussion

Overall, our results highlight the advantages of the EDA framework for biological systems with intermittent events of interest, even those operating at dramatically different timescales (seconds versus hours). Imaging of mitochondrial divisions benefits from extended imaging duration to increase the odds of capturing such rare events, and fast imaging speed once constrictions form to capture the structural intermediates of membrane remodelling. For *C. crescentus*, EDA allows us to follow the entire cell-cycle and image the final states of division at high temporal resolution. In both systems, as observed from the measured constriction diameters, EDA data gathered from individual cells contained smaller constriction sizes, difficult to capture by traditional imaging approaches due to their transience. Beyond that, EDA generates data with higher event-specific content density: more frames are dedicated to active time periods enriched in events of interest.

The EDA framework is generalizable to other microscopes capable of on-the-fly adaptation of acquisition parameters. For users seeking to implement it, we provide a Micro-Manager plugin (Supplementary Note 1), to assist in its adaptation to different imaging methods and biological systems. The plugin is compatible with multithread processing based on a user-provided neural network; thus, with sufficient computational power the response time of the adaptive feedback is limited by the time to process a single frame. While our implementation adapts the imaging speed in a discrete manner, EDA could evoke more complex responses such as multi-level or continuous actuation of the frame rate, as well as controlling other acquisition parameters such as the excitation power or exposure time. The benefits of applying EDA will depend on the specific dynamics of the observed events the greatest advantages are expected if the sample and microscope are amenable to a large difference in imaging speeds. Brief, dynamic and rare events that can be predicted in advance are therefore best suited for observation using EDA.

As with all adaptive microscopies, care should be taken to decouple variations in experimental parameters from real biological changes. For example, phototoxicity varies non-linearly with the light dose^40^, and individual experiments using EDA will differ in the cumulative light dose based on the event-specific content of the sample. Consider the case of altered genetic backgrounds that change the frequency of events of interest: EDA would result in a different mean cumulative light dose for measurements of the same duration. Therefore, events collected under similar doses should be compared, and large sample sizes and randomization will be beneficial when pooling datasets acquired using EDA.

Our work contributes to the developing field of ‘smart’ or ‘self-driving’ microscopes^2,41^. By adapting acquisitions on-the-fly, microscopes can protect a sample from unnecessary illumination^9–16^, generate faster^42,43^ or better quality data^44–46^, optically highlight cells for genetic screening^23^, pinpoint single molecules with vanishingly small numbers of photons^47^, or manipulate nanoscale objects^48^. On the other hand, machine learning has proven useful for many image analysis tasks, allowing the detection of increasingly complex sample features. As event recognition networks become increasingly accessible, EDA can enhance the relevance of gathered data by repurposing such networks to actuate changes in acquisition routines. The synergy between nuanced detection and multifaceted adaptation offers a powerful means to boost biological discovery.

## Methods

### Sample preparation

#### Cos-7 cells for mitochondrial imaging

African green monkey kidney (Cos-7) cells were obtained from HPA culture collections (COS7-ECACC-87021302) and were cultured in Dulbecco’s modified Eagle medium (DMEM, ThermoFisher Scientific, 31966021) supplemented with 10 % fetal bovine serum (FBS) at 37°C and 5 % CO_2_. For imaging, cells were plated on 25 mm, #1.5 glass coverslips (Menzel) 16-24 h prior to transfection at a confluency of ∼1×10^5^ cells per well. Lipofectamine 2000 (ThermoFisher Scientific, 11668030) was used for dual transfections of Cox8-TagRFP and Emerald-Drp1. Transfections were performed 16-24 h before imaging, using 150 ng of plasmid and 1.5 µl of Lipofectamine 2000 per 100 µl Opti-MEM (ThermoFisher Scientific, 31985062). Imaging was performed at 37°C in pre-warmed Leibovitz medium (ThermoFisher Scientific, 21083027).

#### Caulobacter crescentus cells for bacterial imaging

Liquid *C. crescentus* cultures were grown overnight at 30 °C in 3 ml of 2x PYE^49^ (Peptone, Merck, #82303; Yeast Extract, Merck, #Y1626) medium under mechanical agitation (200 rpm). Liquid cultures were re-inoculated into fresh 2x PYE medium to grow cells until log-phase (OD_660_=0.2-0.4). Antibiotics (5 µg ml^−1^ kanamycin and 1 µg ml^−1^ gentamicin) were added in liquid cultures for selecting cells containing fluorescent protein fusions. To induce the expression of FtsZ-sfGFP and mScarlet-I under the *P*_*xyl*_ and *P*_*van*_ promoter respectively, 0.5 mM of vanillate and 0.2 % weight/volume xylose were added to the culture 2 hours before imaging or synchronization^50^.

*C. crescentus* cell cultures were spotted onto a 2x PYE agarose pad for imaging. To make the agarose pad, a gasket (Invitrogen™, Secure-Seal™ Spacer, S24736) was placed on a rectangular glass slide, and filled with 1.5 % 2x PYE agarose (Invitrogen™, UltraPure™ Agarose, 16500100) containing 0.5 mM of vanillate and 0.2 % of xylose. No antibiotics were added. Another glass slide was placed on the top of the silicone gasket, and the sandwich-like pad was placed at 4 °C until the agarose solidified. After 20 min, the top cover slide was removed, and a 1-2 µl drop of cell suspension was placed on the pad. After full absorption of the droplet, the pad was sealed with a plasma-cleaned #1.5 round coverslip of a diameter of 25 mm (Menzel). For imaging the synchronized cells, a 1x PYE agarose pad was used to let cells grow slower.

### iSIM imaging

Imaging was performed on a custom built instant structured illumination microscope (iSIM), previously described in detail^25^. Emerald/sfGFP and TagRFP/mScarlet-I fluorescence was excited using 488 and 561 nm lasers respectively. A system of microlens and pinhole arrays together with a galvo actuated mirror allows for instant super-resolution imaging on the camera chip (Photometrics Prime95B sCMOS). The implementation of a mfFIFI setup allows for homogeneous excitation over the full field of view^25^.

### Event detection

#### Construction of input and output datasets

Detection of potential division events was performed using a neural network with U-net architecture^38^. An input dataset was constructed using 3700 dual-color images labeling mitochondria and Drp1, 1000 of which contain states close to division. This dataset was enhanced 10-fold by rotating the images, to make up the final dataset (37000 images). This data set was recorded on a fast dual-color SIM setup at Janelia Research Campus^51^ and published previously^17^, but processed via Gaussian filtering to imitate the expected resolution of raw iSIM data before deconvolution. The output ground truth dataset was generated from the fluorescence channels, followed by manual curation. Briefly, to identify division sites in the presence of a molecular marker, we identified constriction sites by looking for local saddle points. Saddle points are characterized by opposing principal curvatures, or eigenvalues of the local Hessian matrix. The product of the principal curvatures, or the Gaussian curvature, will therefore be negative - as will be reflected by the determinant of the local Hessian matrix. Therefore, computing the Hessian matrix for both the mitochondrial and the Drp1 channel highlights saddle-points in the mitochondrial channel overlapping with high-intensity Drp1 spots.

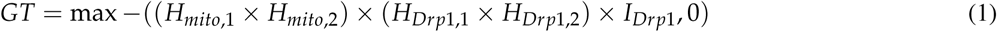

where *H*_*channel,i*_ represents the largest or smallest eigenvalue of the Hessian matrix for *i* = 1, 2 respectively, *I*_*channel*_ represents the raw fluorescence channel and *×* element-wise multiplication. The final *GT* matrix is normalized so that the full range is rescaled to 0-255 to make 8-bit heat-maps, with higher values representing more likely constriction sites.

Since mitochondrial division sites represent a large fraction, but not all events with Drp1 overlapping with mitochondrial saddle points, the processed frames were then curated manually to increase accuracy for real constriction events by visually removing false positives or ensuring consistent detection of active constriction sites. False positives most frequently corresponded to close mitochondrial contacts with nearby Drp1 bound to the outer membrane (but not constricting), very bright Drp1 spots and mitochondria with bi-concave disk-like shapes where thinning of the mitochondrion gets mistaken for a constriction site. False negatives were most often due to very late stages of fission or moments after fission as well as the onset of constriction which is nuanced and develops progressively. This ensured that the event detection network was able to discriminate events of interest with high accuracy. The dataset was then split 80:20 into training (29600 images) and testing (7400 images) datasets.

#### Network architecture and training

The network was trained on 29600 dual-color images (128×128×2) of mitochondria and Drp1 as input, and using the groundtruth images (128×128) marking the respective locations of divisions as output (Supplementary Note 2). We used pixel-wise mean-squared error as loss function, and trained the model using the Adam optimizer for 20 epochs with a batch size of 256. As neural network architecture we used a U-Net^38^ of depth 2 and kernelsize 7×7, with the number of initial feature channels set to 16 that were doubled after every pooling layer. Training was performed using tensorflow/keras and python 3.9.^52^.

#### Assessment of network prediction accuracy

To assess the accuracy of the network, individual event scores were first isolated from the heat maps using an intensity threshold of 80, since weak predictions and weak ground truth signals represent a low relative event probability and as such, will not contribute to the decision making step of the acquisition. Then, instead of counting discrete events, each event was weighed by its ground truth or predicted intensity: true positives were weighed by normalized ground truth intensity (representing what was the true value of the event that was detected), false positives were weighed by normalized predicted intensity (representing how wrong the prediction was, given no event was present) and false negatives were weighed by normalized ground truth intensity (representing the cost of missing the ground truth event). This way the strength of the prediction and true underlying signal is taken into account when evaluating the performance of the network. The total score for true positives was then divided by the sum of total true positives (false positives and false negatives) to produce an accuracy metric.

With this, the network reaches an accuracy of 86.7% when tested on test data, that was not used during training, with 5.4% false positives and 7.9% false negatives. The output of inference is a two dimensional map of relative division probabilities in the range 0 to 255. Inspecting false positives shows that events that most frequently contributed to false positives were close mitochondrial contacts with nearby Drp1 bound to the outer membrane (but not constricting), very bright Drp1 spots and mitochondria with bi-concave-disk-like shapes where thinning of the mitochondrion gets mistaken for a constriction site. False negatives were most often due to very late stages of fission or moments after fission as well as the onset of constriction which is nuanced and develops progressively.

### Event-driven acquisition for adaptive temporal sampling

The different parts of the EDA framework were implemented in separated modules that allowed for continuous and independent testing of the components.

#### Data handling

The EDA framework is distributed over Micro-Manager^20^ for general microscope handling, and Matlab (MathWorks) for the control of the timing of the microscope components and Python for network inference. Furthermore, the Python module was used on a machine in the network due to hardware restrictions on the local computer used for microscope control. A network attached storage (NAS) unit was used to allow for communication of the different components of the system over the local 10 Gbit network. Recorded frames were stored as single .tif files to the NAS by Micro-Manager. A server implemented using the watchdog module (Python) on the remote machine detected new files for inference. After calculation of the decision parameter, the value was saved to a binary file that was used by Matlab to calculate the values for the next sequence of imaging.

#### Event Server

When a new file for each channel respectively was detected on the NAS, the event server implemented in Python running on a remote machine first performed data preparation on the recorded frames. The frames were resized by a factor of 0.7 to match the pixel size of the training data. Both frames were smoothed using a Gaussian with *σ* 1.5 px with an additional background subtraction in the Drp1 channel (Gaussian with *σ* 7.5 px). The frames were tiled into overlapping 128×128 px sized tiles to match the size of the training data. The individual tiles of the structure (mitochondria/*C. crescentus*) channel were again normalized individually.

Inference was performed on pairs of structure/foci tiles and the output was stitched together. The maximum value from the stitched frame was recorded as the highest relative probability in the frame for a division event and saved to the binary file on the NAS.

#### Hardware control

The timing of the microscope hardware is controlled using a PCI 6733 analog output device (National Instruments). The device is used in background mode requesting data when the existing sequence buffer has 1 s remaining. For mitochondria imaging, the slow mode sends a sequence containing one frame over five seconds (0.2 Hz) and a sequence containing five frames in one second (5 Hz) in the fast mode. For bacterial imaging, sequences with one frame over 3/9 or 2/12 minutes is provided for slow/fast and normal or synchronized colonies respectively. The parameters are chosen depending on which value was read from the binary file on the NAS that contains the latest event information written by the event server. In addition to a threshold, a hysteresis band is implemented here by defining an upper and a lower threshold. Fast acquisition starts at surpassing the upper threshold and is only stopped when the event score falls below the lower threshold. For *C. crescentus* imaging, the number of fast frames was further set to a minimum of three.

Due to the fast imaging rates in the mitochondria imaging, the new sequence is calculated before the event server has calculated the relative probability map for the last frames, leading to a delay in the reaction of the EDA framework. This is overcome for the *C. crescentus* imaging by delaying the calculation of the new sequences by 10 seconds allowing for mode switching on the newest data available.

### Data Analysis

Statistical significance of differences in averages reported by asterisks was calculated using the the independent two-sample t-test for minimal constriction widths (Figures 3 & 4) and the Welch’s t-test for event scores (Suppl. Note 2) and number of events (Figure 3). Equality of variances in the data sets to decide between the two was tested for by the Levene test. All implementations were used as provided by the scipy.stats Python package.

#### Photobleaching Decay

The different photobleaching kinetics of the modes were characterized by the intensity contrast of the samples. The channel of the structural feature (mitochondria and caulobacter outline) was segmented using a Otsu-thresholding method after a median and Gaussian filter were applied (kernel size of 5 px each). The intensity contrast was then calculated as the mean intensity in the segmented region divided by the mean intensity in the rest of the image. Variation in the starting contrast between series was accounted for by normalization of the starting intensity contrast values. The decay constant was then obtained by fitting an exponential decay function (*y* = *a* exp^(−*bx*)^ +*c*) to the intensity contrast over time data and extracting the *b* term. The reported imaging time ratios were calculated from the times from start of imaging to 90 % of intensity contrast for each series.

#### Cumulative Light Dose

The cumulative light dose over time was taken to be proportional to the number of previously recorded frames. This is appropriate here, since the iSIM has a constant scan speed, and we did not modulate the laser intensity. Therefore, each sample received a fixed light dose per frame. Experiments were truncated after a user-defined minimum intensity contrast (SNR) was reached (1.1 for mitochondria and 1.02 for *C. crescentus*).

### EDA Event Evaluation

The number of frames per event was calculated by counting the number of frames recorded for a fixed time after EDA triggered fast imaging or would have triggered fast imaging in slow mode. Events of interest were defined by a minimum value of the neural network output of 80 (90 for *C. crescentus*). Frames were analyzed until a maximal observation time of 20 seconds (1 hour for *C. crescentus*) was reached.

#### Constriction Width

The slow and EDA imaging modes were compared by calculating the minimal width of constriction measured during an event as described above. The measurement method was similar to that described in^17^. The deconvolution of the images was performed using the Richardson Lucy algorithm as implemented by the flowdec Python package with 30 iterations^53^. Segmentation, skeletonization and spline fitting the resulting points led to a backbone of the mitochondrion in a frame of 20 × 20 pixels around the position of the detected event. 100 perpendicular lines were generated around the closest point of the backbone to the event position. The intensity profile along those lines was fitted using a Gaussian profile. The full width at half maximum (FWHM) of the Gaussian profile with the smallest *σ* was recorded as the measured width for the frame. The minimal width measured for an event was calculated as the minimal FWHM over the observation time times the pixel size of the iSIM setup (56 nm).

## Code and Data Availability

All code used in this project is available at https://github.com/LEB-EPFL/EDA. The data contained in this manuscript and the training data for the model used can be found at 10.5281/zenodo.5548354. The Python plugin described can be found at https://github.com/wl-stepp/eda_plugin.

## Acknowledgements

We thank Hélène Perreten for technical support with cell culture and plasmid construction, and Laurent Casini and Justine Collier (University of Lausanne) for sharing plasmids and protocols for the bacterial experiments. Imaging data used for training the neural network in this publication were produced in collaboration with the Advanced Imaging Center (AIC), a facility jointly supported by the Gordon and Betty Moore Foundation and HHMI at HHMI’s Janelia Research Campus. We thank Lin Shao and Teng-Leong Chew at Janelia AIC for their help with SIM imaging.

This work was supported by the Swiss National Science Foundation project grant (SNSF; 182429 to S.M., D.M. and W.L.S.), and National Centre for Competence in Research (NCCR Chemical Biology; S.M. and D.M.); and the European Union’s H2020 programme under the the European Research Council (ERC; CoG 819823 Piko, S.M. and C.Z.), and the Marie Skłodowska-Curie Fellowships (BALTIC, J.G.). MW is supported by the EPFL School of Life Sciences and a generous foundation represented by CARIGEST SA.

## Author contributions

D.M., J.G. and S.M. conceived and designed the project. D.M., M.W. and S.M. supervised the project. D.M. collected the training data and performed the experiments on mitochondrial division. D.M. and M.W. implemented the neural network for event detection. W.L.S. and D.M. implemented the EDA framework and performed data analysis. W.L.S. developed the Python plugin for Micro-Manager and its documentation. W.L.S performed the experiments on *C. crescentus*. C.Z. cultivated the *C. crescentus* strains and prepared samples for imaging. W.L.S prepared the figures. D.M., W.L.S. and S.M. wrote the manuscript with contributions from all authors.

