## Supplementary Information for "Event-driven acquisition for content-enriched microscopy"

### Supplementary Note 1: Micro-Manager Plugin

Our original implementation of event-driven acquisition for imaging speed adjustment was specific to our custom-built iSIM microscope. To extend its use for other setups, we developed a Python-based plugin for Micro-Manager. The plugin allows for modular combination of different components, to allow others to adapt the functionality of the implementation presented in this manuscript. The main additional capabilities are control of fully Micro-Manager driven microscopes with standard and Micro-Magellan/Pycro-Manager[54,55] driven acquisitions. The event detection using neural networks presented here can be combined with the different control mechanisms owing to the modular architecture of the plugin.

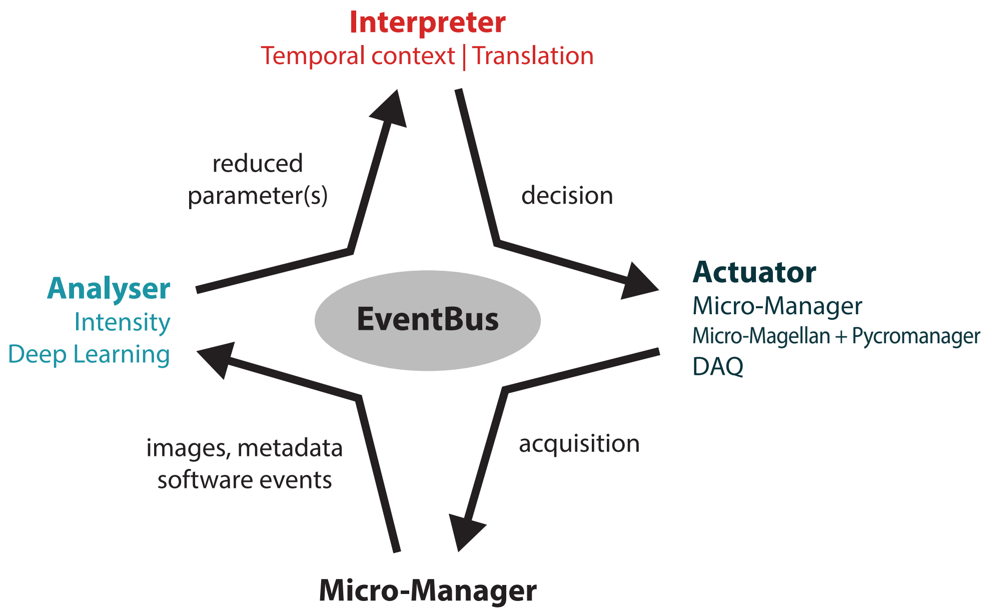


Information gathered by Micro-Manager is transferred to an **analyser**. It gathers the amount of images/data it needs to perform analysis and handles resource allocation, as this step will typically be the bottleneck for the EDA loop. This results in a reduced set of parameters (event score/probability) which is reported to an interpreter. The **interpreter** gathers the information from different analysis threads and keeps track of the temporal context. Depending on this data and parameters set by the user it decides for an action to be taken by an actuator. The **actuator** changes the parameters of the acquisition to implement the change.

With the implementations of these modules, many standard microscopes can be equipped with EDA. The use of a custom network that is based on tensorflow/keras is supported for 2D/3D and multiple channel models. The implementation of a hardware synchronized actuator shows a possible implementation of EDA for a custom setup.

More information can be found here:
https://github.com/wl-stepp/eda_plugin
https://event-driven-acquisition.readthedocs.io/

### Supplementary Note 2: U-Net for the detection of divisions

The U-Net used for event detection in this project was trained with images of both DRP1 and the mitochondria outline. The ground truth was obtained as discussed in the Methods. Scale bars: 1 μm.

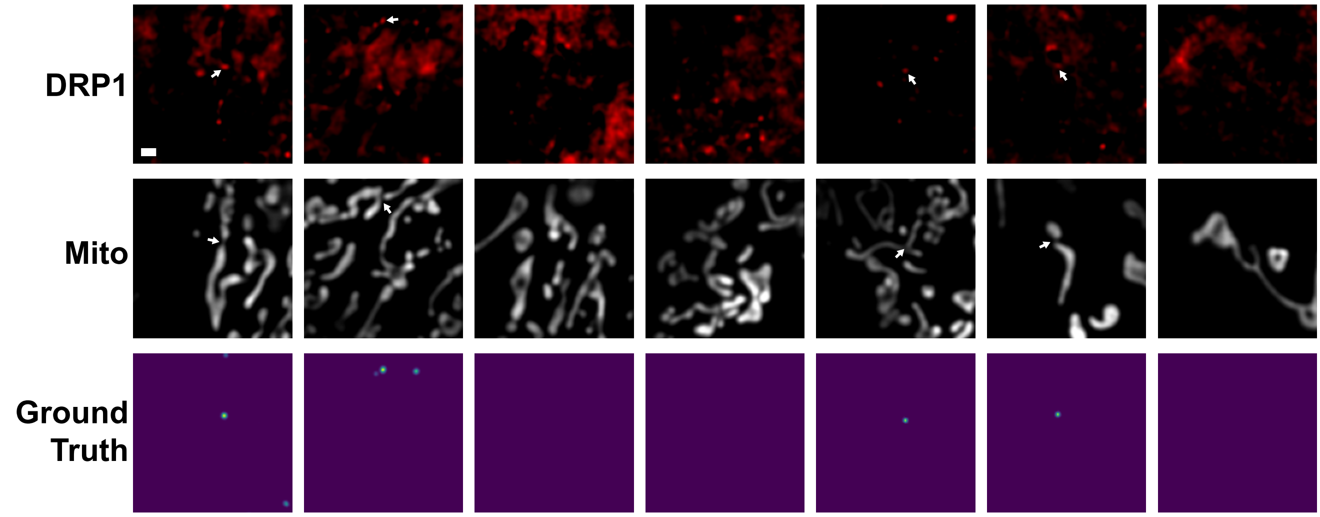


The resulting network also gives reliable predictions of division precursor states when applied to data from the instant SIM used for the implementation of EDA. Frames here are not deconvolved to represent the data as supplied to the network. Using the super-resolved images directly from the iSIM without deconvolution allows for faster reaction times of the system.

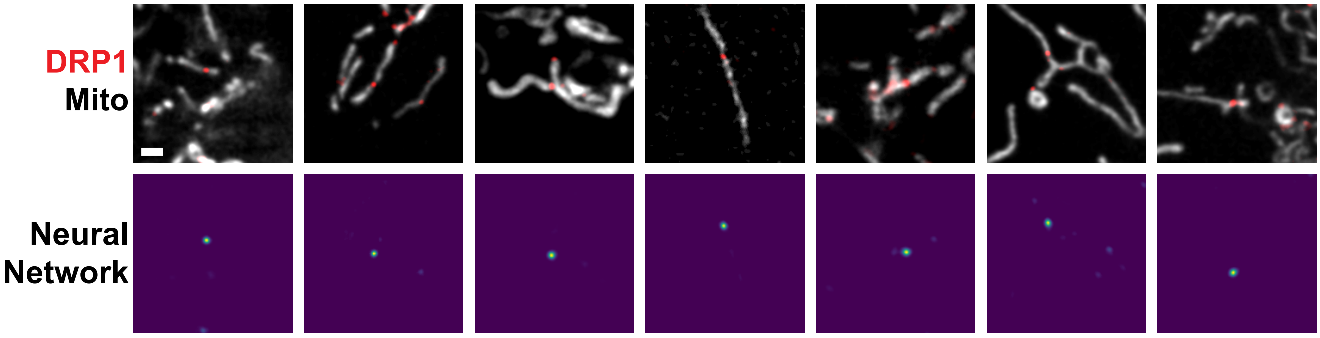


Due to morphological similarities and the presence of a constriction marker in FtsZ, the same network without additional or separate training also marks constriction sites in *C. crescentus* cells that are close to division. States just after division do not show high event scores (Images 2 and 3) as well as cells with a punctate but polar FtsZ configuration, showing the benefit of including both channels in the neural network analysis.

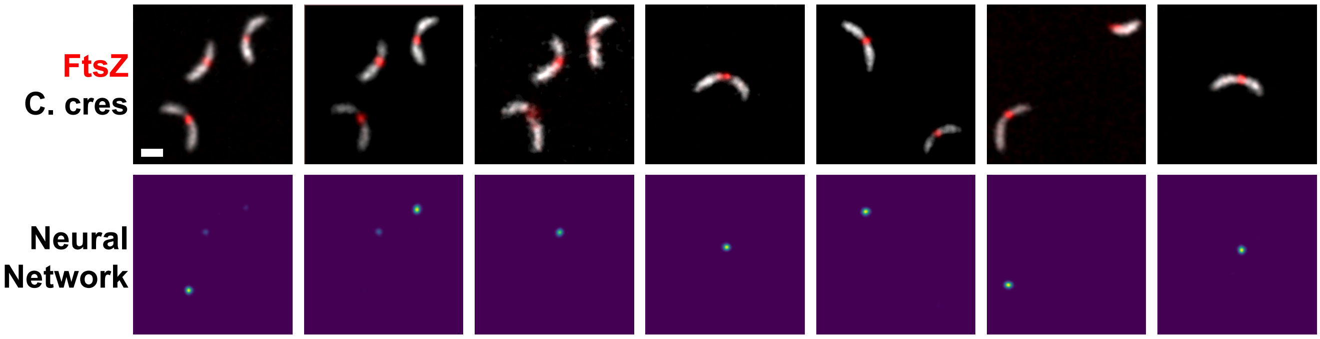

The network's ability to trigger EDA was examined by extracting the total event scores for each frame for EDA and traditional imaging approaches, as well as separating the data in EDA experiments into the frames obtained at slow imaging speed (EDA-slow) and fast imaging speed (EDA-fast). The event score is higher for both fast data subsets, showing that the network correctly triggered the fast speed. Moreover, the events are long and consistent enough, and EDA reaction times are fast enough to image events with a high score with the higher imaging speed. As more data is collected at higher event-scores, the overall event score in EDA experiments is also higher compared to fixed imaging speed experiments.

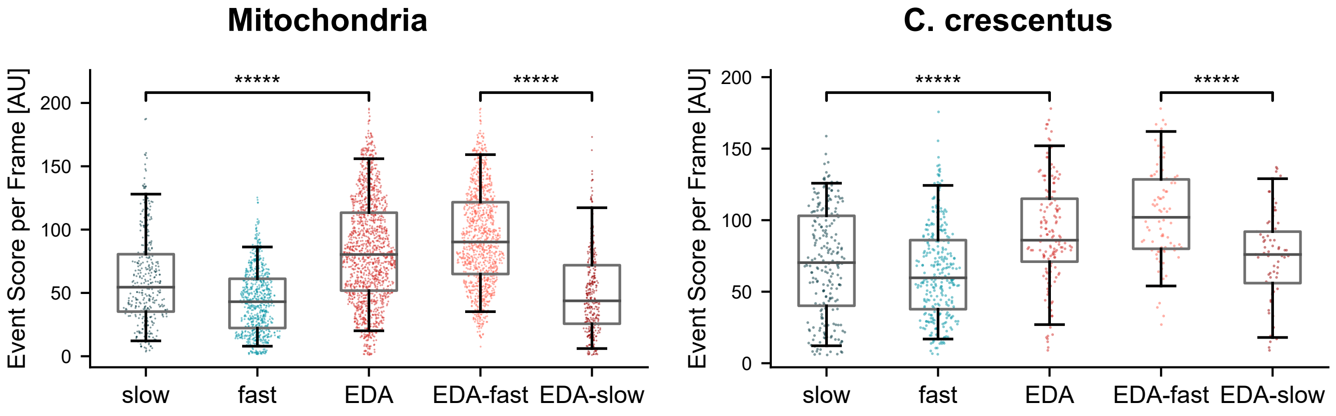


From the training datasets, we identified events that had a high putative ground truth, but were not associated with precursors to division (false positives). The events that most frequently contributed to false positives were close mitochondrial contacts with nearby DRP1 bound to the outer membrane (but not constricting), very bright DRP1 spots and mitochondria with biconcave-disk-like shapes where thinning of the mitochondrion gets mistaken for a constriction site. We also observed cases of division that were not associated with a high putative ground truth (false negatives). False negatives were most often due to very late stages of fission or moments after fission as well as the onset of constriction which is nuanced and develops progressively. Through manual curation, we improved the network's ability to avoid assigning these a high event score.

### Examples of features requiring manual curation

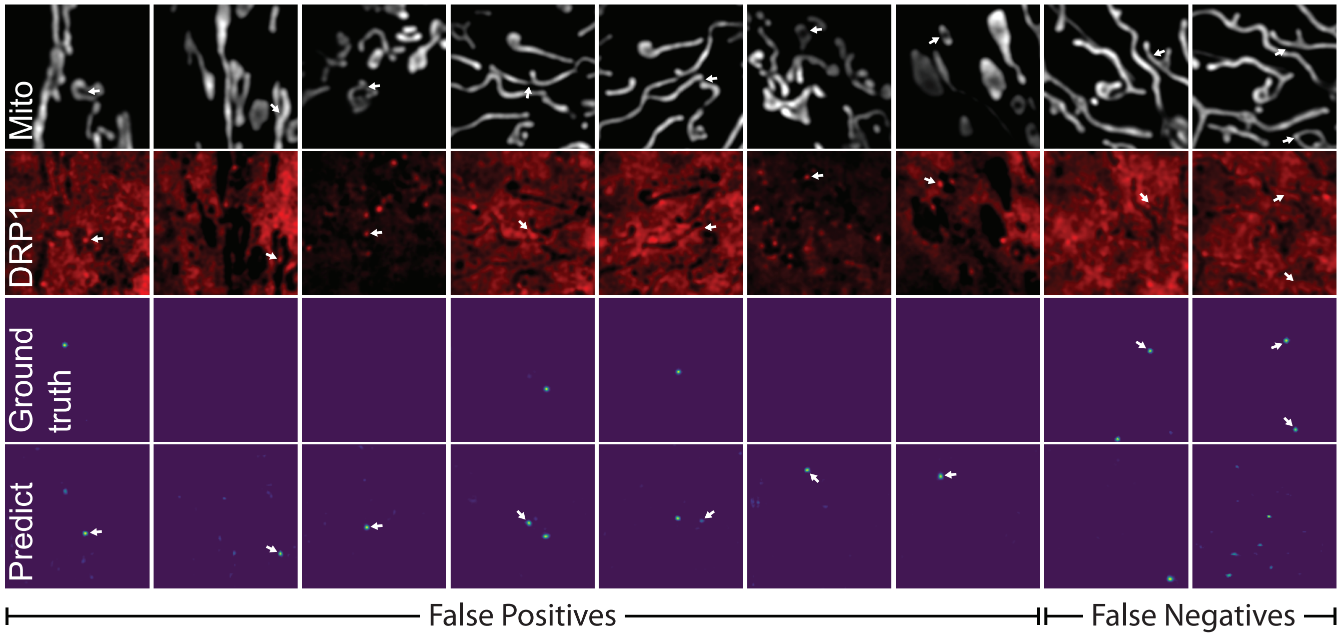

### Supplementary Note 3: Strains and plasmids

#### *Caulobacter crescentus*

The strains and plasmids used for this study are summarized in Table 1 below. The wild-type strain (CB15N) was electroporated with the $P_{xyl}$::ftsZ-sfGFP plasmid and the *PMT335-mScarlet-I* plasmid sequentially to yield the dual-color strain. Notably, the DNA sequence of sfGFP and mScarlet-I used for this study was optimized for respective protein expression in *C. crescentus*. The two plasmids have been deposited at Addgene with the ID 174505 ($P_{xyl}$::ftsZ-sfGFP) and 174506 (*PMT335-mScarlet-I*).

| **Resources** | **Description** | **Source** |
|---|---|---|
| **Strains** | | |
| CB15N | NA1000, synchronizable derivative of wild-type CB15 | (Lambert et al, 2018)[56] |
| CB15N $P_{xyl}$*::ftsZ-sfGFP* $P_{van}$*::mScarlet-I* | CB15N electroporated by the $P_{xyl}$*::ftsZ-sfGFP* and *PMT335-mScarlet-I* plasmids sequentially | This study |
| **Plasmids** | | |
| $P_{xyl}$*::GFPC-2* | Integrated plasmid for protein expression in *C. crescentus* under xylose induction, kanamycin resistant | (Lambert et al, 2018)[56] |
| $P_{xyl}$*::ftsZ-sfGFP* | Plasmid harboring *ftsZ-sfGFP* gene which replaces the *GFP* in the $P_{xyl}$*::GFPC-2* plasmid | This study |
| *pMT335* | High-copy number plasmid for protein expression in *C. crescentus* under vanillate induction, gentamicin resistant | (Thanbichler et al, 2007)[57] |
| *PMT335-mScarlet-I* | *PMT335* plasmid harboring the *mScarlet-I* gene for its cytoplasmic expression | This study |

**Table 1:** Strains and plasmids used for this study

### Supplementary Note 4: Timing

We performed tests of the performance of the EDA analysis pipeline. Both the original distributed implementation and the Micro-Manager plugin were tested on 1024x1024 images. For both implementations, the analysis pipeline was running on a PC with an Intel Xeon E5-2620, 16 GB RAM, a NVIDIA GeForce GTX 1080 Ti and 10 Gbps connectivity.

#### A. Distributed Implementation

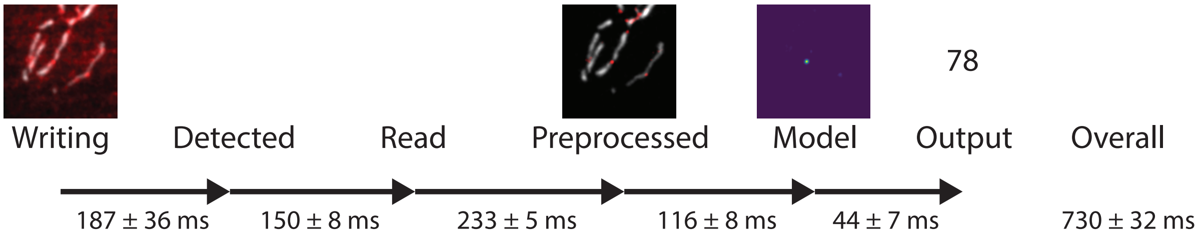


For the distributed implementation, Micro-Manager was running on a PC with an Intel Xeon Silver 4210R CPU, 128 GB RAM and 10 Gbps connectivity to the network attached storage. In this implementation, a considerable amount of time is spent on the input/output procedures of the images. Detection and reading of the data from the network attached storage took around half of the full time of the analysis procedure. The hardware synchronized control of the microscope was set to poll for new data every 1s, so EDA full-circle reaction times would typically lie in between around 0.8 and 1.8 s. In our mitochondria experiments (with 2 channels, 100 ms exposure each), this resulted in 4 to 7 frames with the fast imaging speed before switching to the slow speed. For a switch from the slow imaging speed to the fast imaging speed, we decided to always take another frame with the long interval before switching to the fast speed in order to avoid additional frame delay values in the data. For the experiments on *C. crescentus*, the hardware control was set to check the output value 10 seconds after the acquisition of the images, enabling the switch of imaging speed without delay.

#### B. Micro-Manager Plugin

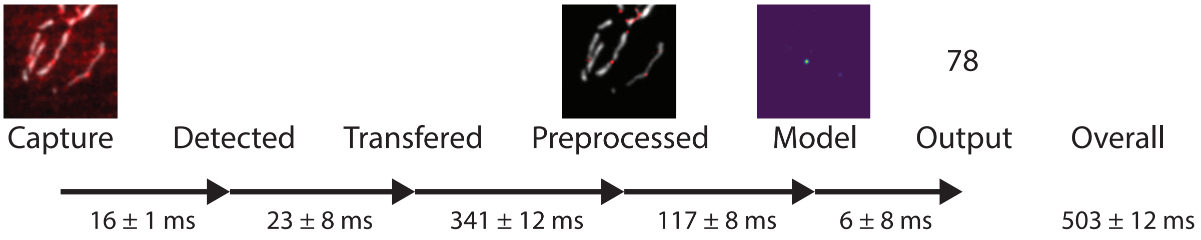


In the implementation of the Micro-Manager plugin we reduced the overhead of data I/O by almost a factor of 10 compared to the distributed implementation. The multithreaded implemenation of the preprocessing described in the documentation could be the cause for the slightly higher time needed for pre-processing the images.

### Supplementary Figure 1: Bleaching behavior during EDA imaging

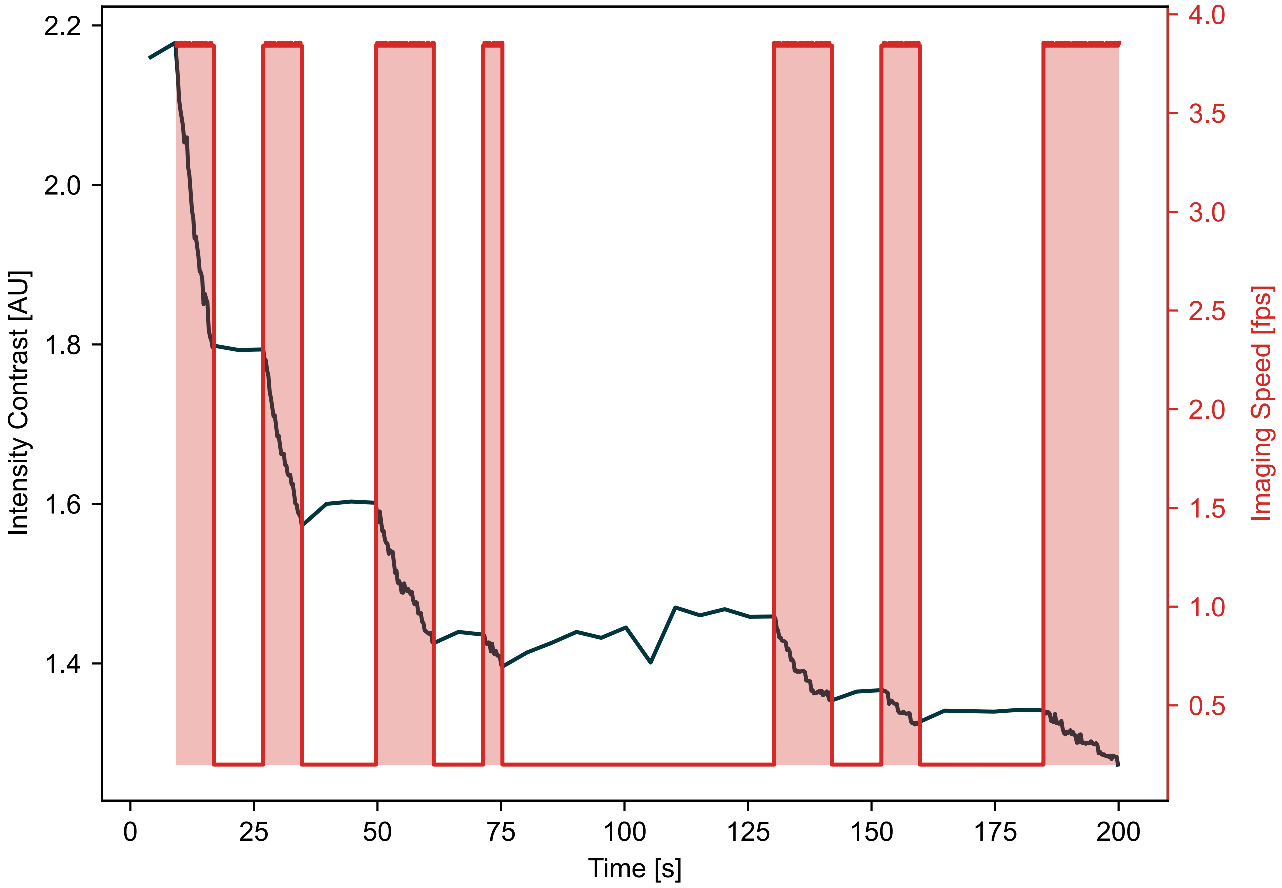


**Supplementary Figure 1: Bleaching behavior of a mitochondria sample during EDA imaging.** The different modes of imaging can clearly be seen in the bleaching curve represented by the signal-to-noise ratio calculated from the intensity inside the mitochondria compared to the signal outside of the mitochondria. For some parts with low frame rate, even a slight recovery of signal can be observed.

### Supplementary Figure 2: Additional event-specific content collected by EDA during fast imaging

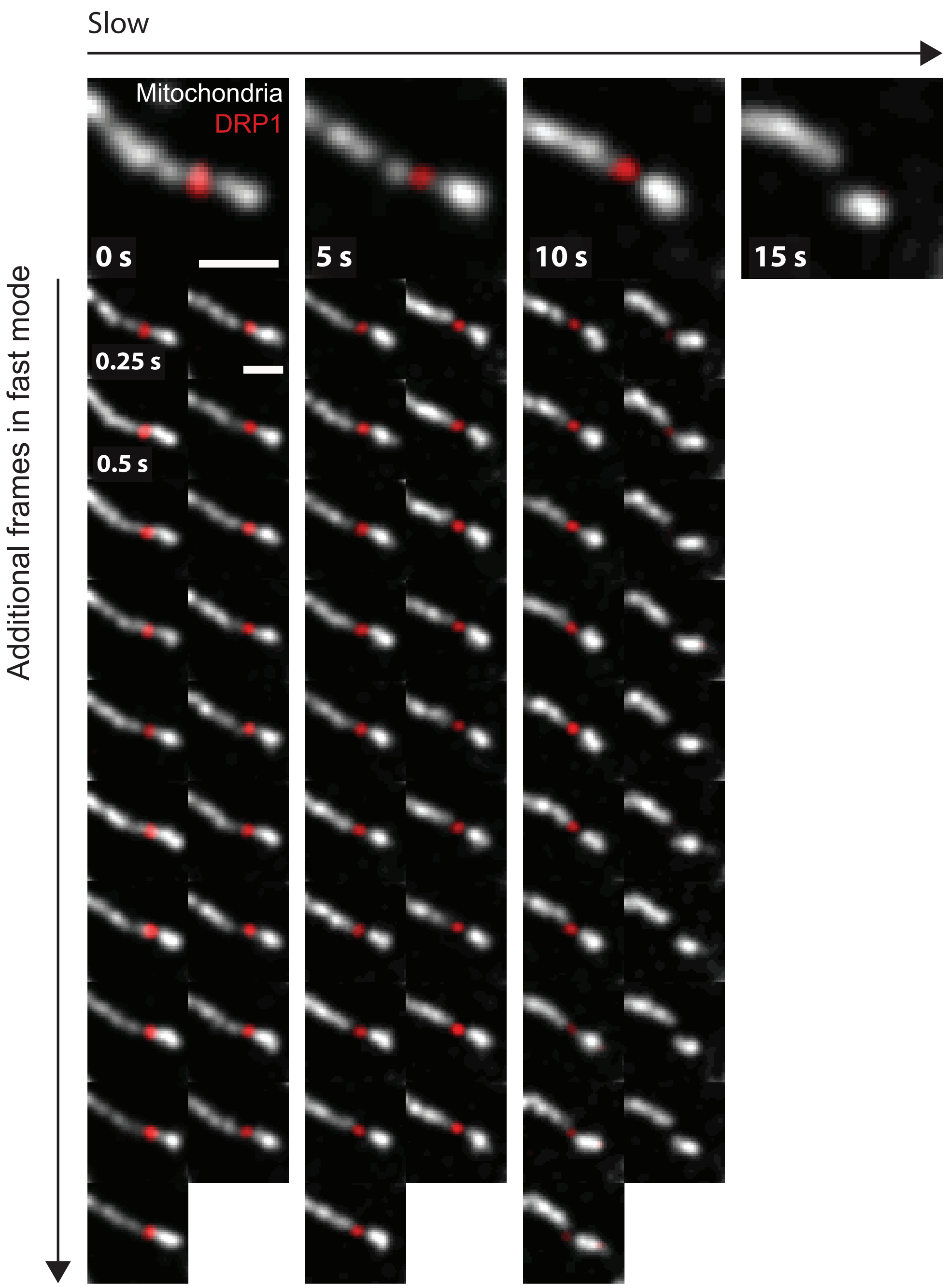


**Supplementary Figure 2: EDA delivers additional frames during events of interest.** Top row: mitochondrial division as it would have been recorded with the slow fixed imaging rate without EDA. Vertical frames: additional frames captured thanks to EDA switching to the fast imaging speed showing more detail of the dynamics of the event. Both the final constriction state and the fade of the DRP1 peak can be observed with higher temporal resolution, enhancing the relevant content of the dataset. This division event can also be seen in Supplementary Video 3.0. Scale bars: 1 µm

### Supplementary Figure 3: EDA for synchronized bacterial imaging

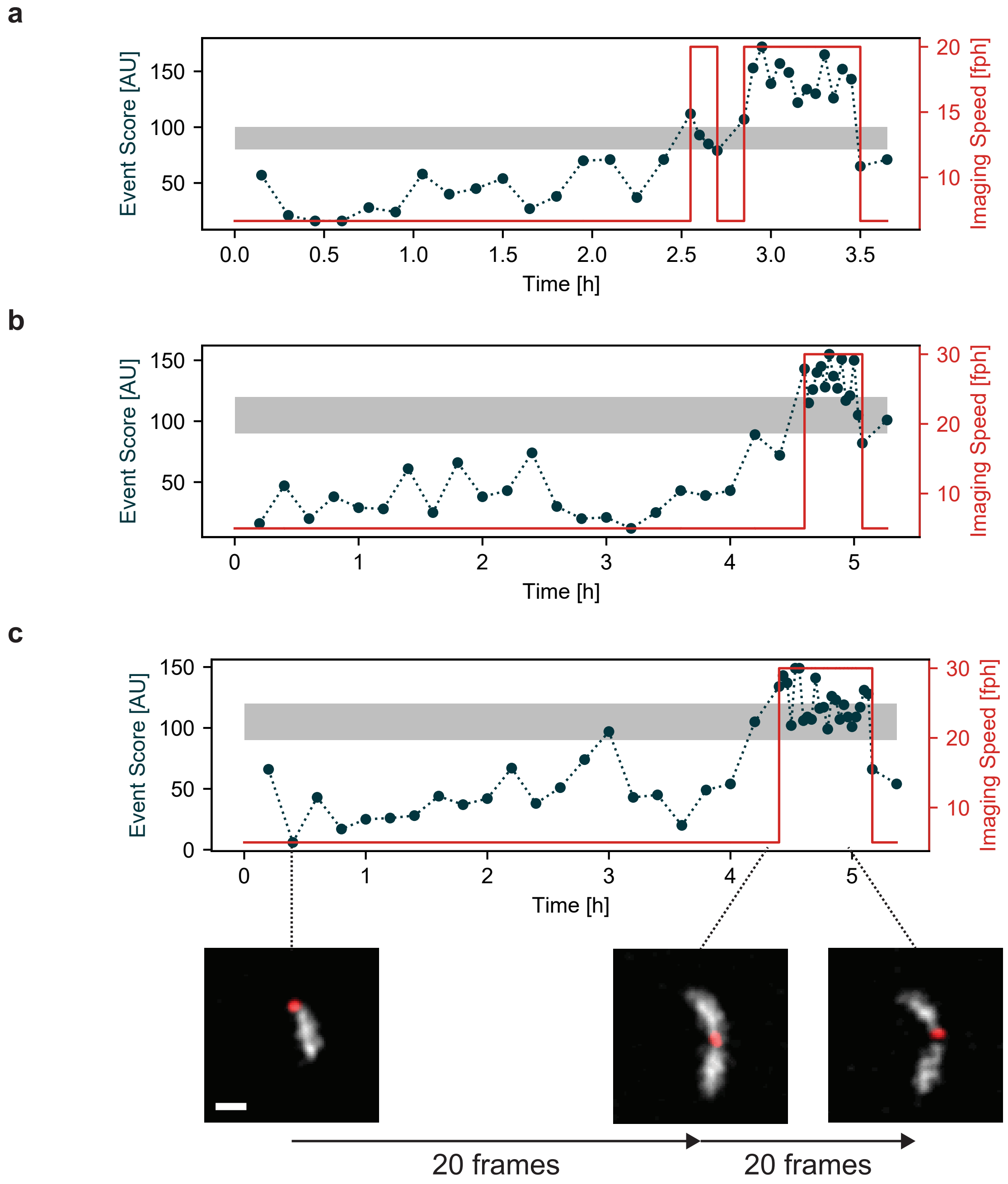


**Supplementary Figure 3: EDA imaging of synchronized bacteria populations.** *C. crescentus*, the strain used in this study, were synchronized via density centrifugation to obtain a population of cells that are all at the beginning of their cell cycle (G0, swarmer). This leads to a time lag before the next divisions take place. As they are synchronized, many bacteria in the sample will then divide at the same time. We used EDA to sense the onset of divisions in the sample and increase imaging speed during the divisions for high SNR and temporal resolution. We tested different times between images for fast and slow speeds, as well as different threshold event scores (grey band). **a** slow: 9 min, fast 3 min. **b and c** slow: 12 min, fast 2 min. Scale bar: 1 µm

### Supplementary Figure 4: EDA highlight extraction 1

The event score output by the neural network can also be used to extract events of high interest from the datasets, after the acquisition is complete. Here, events that triggered EDA in different datasets are shown. The highest event score was used to define a region of interest around the event, representing a time and location of highest interest in the sample.

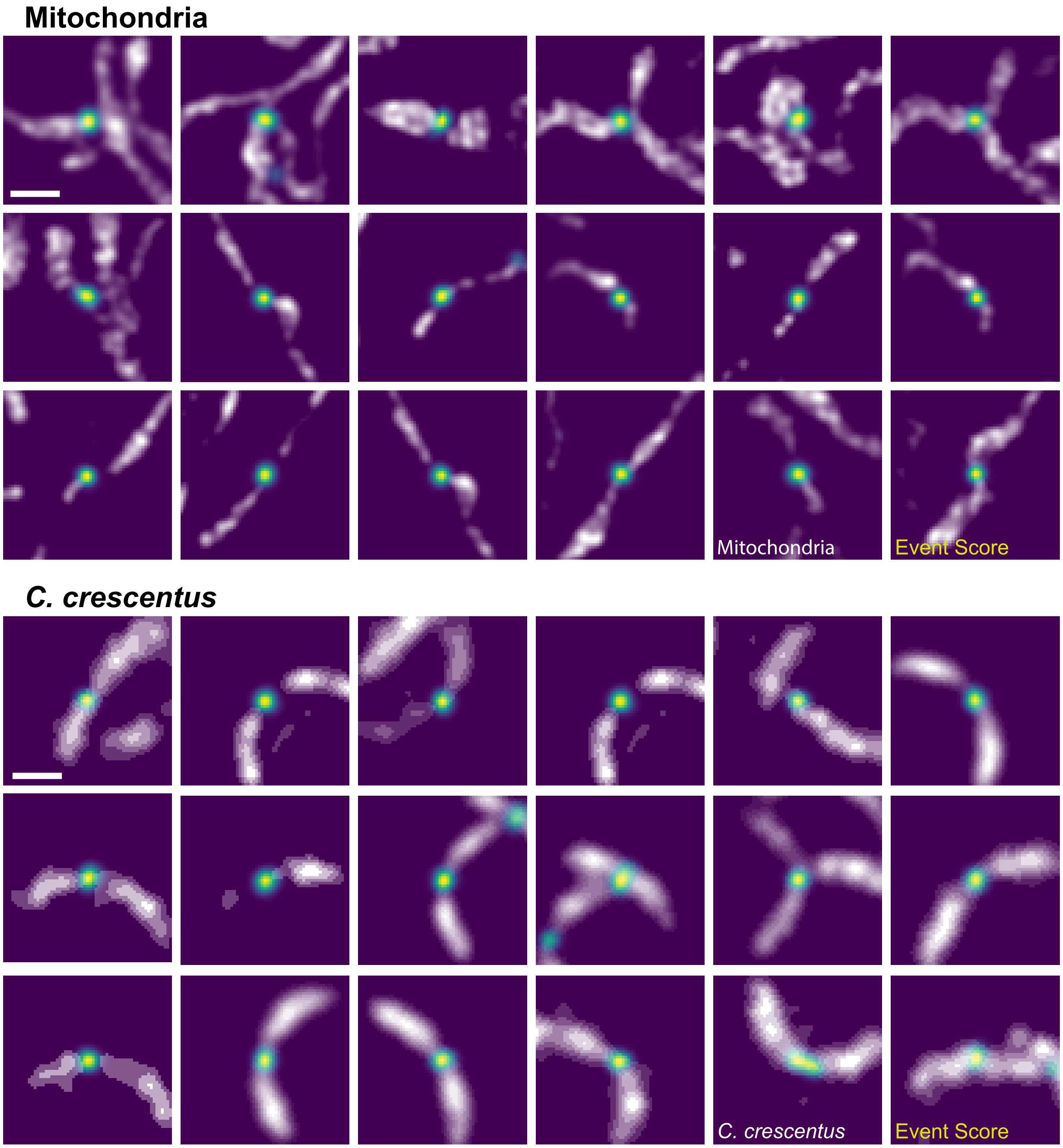


**Supplementary Figure 4: Example maximum event score regions which triggered EDA.** Some regions appear twice, when the neural network event score was high enough to trigger EDA multiple times. Frames are shown in no specific order. Scale bars: 1 µm 4

### Supplementary Note 5: Supplementary Videos

Cropped regions of interest from different EDA experiments with the probability map of the neural network scaled as in Figure 1 (30 to 120).

Real_time: Time per frame proportional to the delay time while recording. Shows the differences in imaging speed. Slow_motion: Time per frame equal for all frames. Shows the higher information content during events of interest.

#### A. Supplementary Video 1.0

Real_time video of a 300x300 px field of view of mitochondria (grey) and DRP1 (red). 3:40 minutes recorded time, fast frame rate 3.8 fps, slow frame rate 0.2 fps.

#### B. Supplementary Video 1.1

Slow_motion movie corresponding to Supplementary Video 1.0.

#### C. Supplementary Video 2.0

Real_time video of a originally 128x128 px field of view of mitochondria (grey) and DRP1 (red). 2:04 minutes recorded time, fast frame rate 3.8 fps, slow frame rate 0.2 fps.

#### D. Supplementary Video 2.1

Slow_motion video corresponding to Supplementary Video 2.0

#### E. Supplementary Video 3.0

Real_time video of a originally 128x128 px field of view of mitochondria (grey) and DRP1 (red). 1:12 minutes recorded time, fast frame rate 3.8 fps, slow frame rate 0.2 fps. Corresponding to Supplementary Figure 2

#### F. Supplementary Video 3.1

Slow_motion video corresponding to Supplementary Video 3.0

#### G. Supplementary Video 4.0

Real_time video of a 856x856 px field of view of *C. crescentus* (grey) and FtsZ (red). 3:54 hours recorded time, fast frames 3 mins, slow frames 9 mins.

#### H. Supplementary Video 4.1

Slow_motion corresponding to Supplementary Video 4.0
